# Isavuconazole therapeutic drug monitoring associated to a pharmacogenetic exploration is far from being a pharmacologists’ whim

**DOI:** 10.1101/316851

**Authors:** L Darnaud, F Lamoureux, C Godet, S Pontier, A Debard, N Venisse, P Martins, D Concordet, P Gandia

## Abstract

Isavuconazole is a new antifungal prodrug to treat invasive aspergillosis and mucormycosis, theoretically not requiring drug monitoring. However, we reported 4 clinical cases with toxic concentrations. Based on Desai’s population pharmacokinetic model, we estimated patients’ kinetic profile. Clearance was abnormally low, likely related to CYP3A4/5 polymorphisms. Thus, we recommend to collect blood sample just before the first maintenance dose to estimate pharmacokinetic profile and individualized dose. For patients presenting high concentrations, pharmacogenetics can be done.

---

Recently, Stott et al. [1] mentioned that “the novel broad-spectrum azole drug isavuconazole does not currently appear to require TDM but ‘real-world’ data are awaited and TDM could be considered in selected clinical cases”. It did not take very long to confirm these proposals and even go further in the recommendations.

Isavuconazole is a new antifungal prodrug approved in the USA and Europe for the treatment of invasive aspergillosis and mucormycosis in adults. The recommended dose is 200 mg (IV or PO) every 8 h for 48 h, followed by 200 mg once daily for maintenance dose applied 12–24 h after the last loading dose. The median trough plasma concentrations of isavuconazole (i.e. value determined just before drug readministration) at steady state (> 7 days of treatment) in patients who received the drug in the SECURE and VITAL trials [2–3], were 3–4 μg/mL. Recently, we observed 4 cases of unexpected high concentrations a few days after the last loading dose, without obvious explaining factors (no drug-drug interactions and no previous hepatic injury). For 2 patients, these high levels were concomitant with adverse effects (patellar tendinopathy and discomfort, respectively).

Isavuconazole plasma concentrations were determined using a validated chromatographic technique according to ISO15189 standards. Using the population pharmacokinetic model published by Desai et al [4], firstly we simulated 50000 pharmacokinetic profiles of isavuconazole for the recommended loading (200 mg/8h x 6) and maintenance doses (200 mg/day). Secondly, for each patient, we computed the Empirical Bayes Estimates (EBE) of the pharmacokinetic parameters (Table 1) and deduced the most likely kinetic profile (Fig. 1-4). Because these patients were extremes with respect to Desai’s model, their EBE’s where computed using an individual parameter distribution with heavier tails than the classical multidimensional log-normal distribution.

**Fig. 1.**
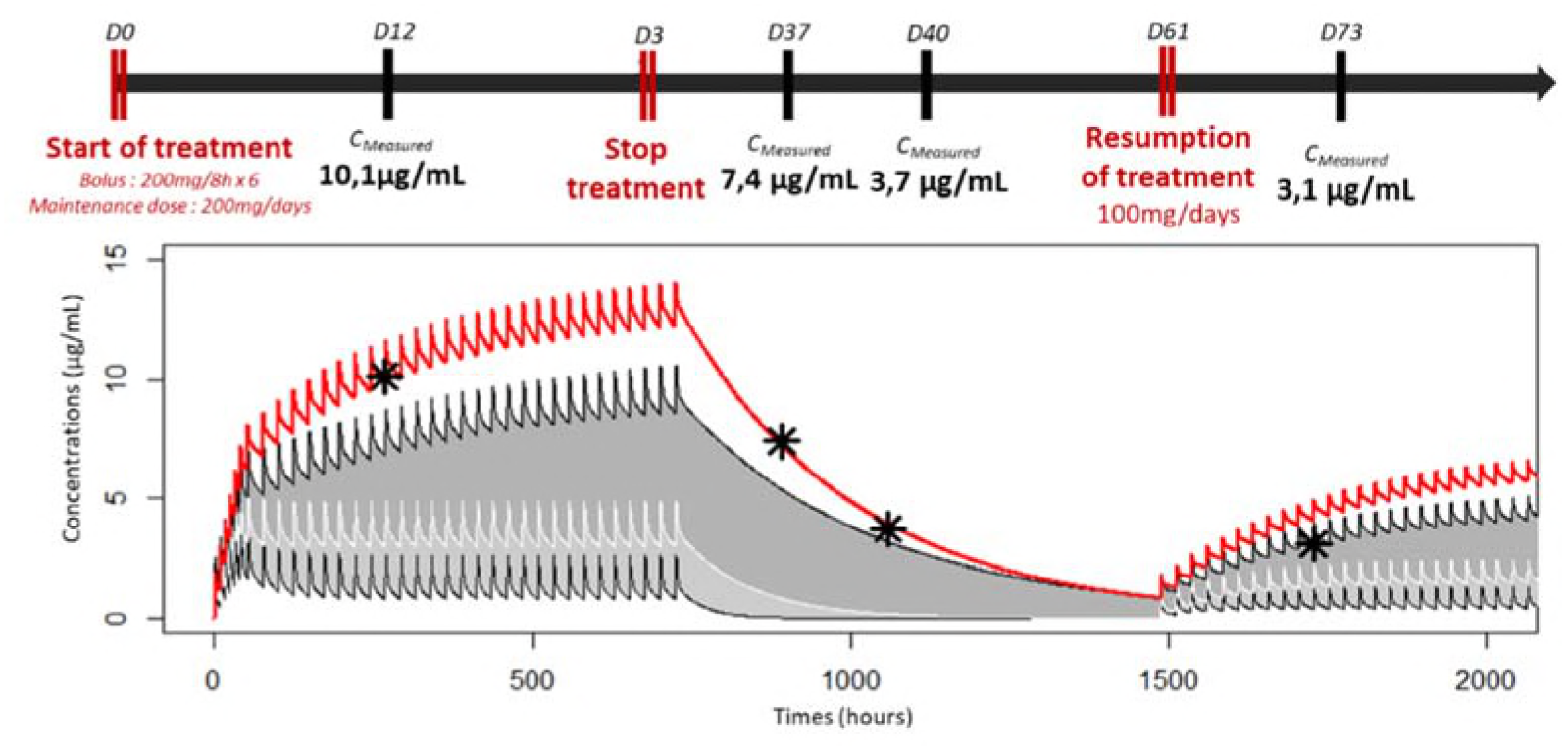
Patient 1: Kinetic profile of isavuconazole. Median kinetic profile (curve in white), “extremes” profiles found in less than 5% and 95% of the population (black curves) and estimated kinetic profile of the patient (red curve); the black stars represent the measured concentrations.

**Fig. 2.**
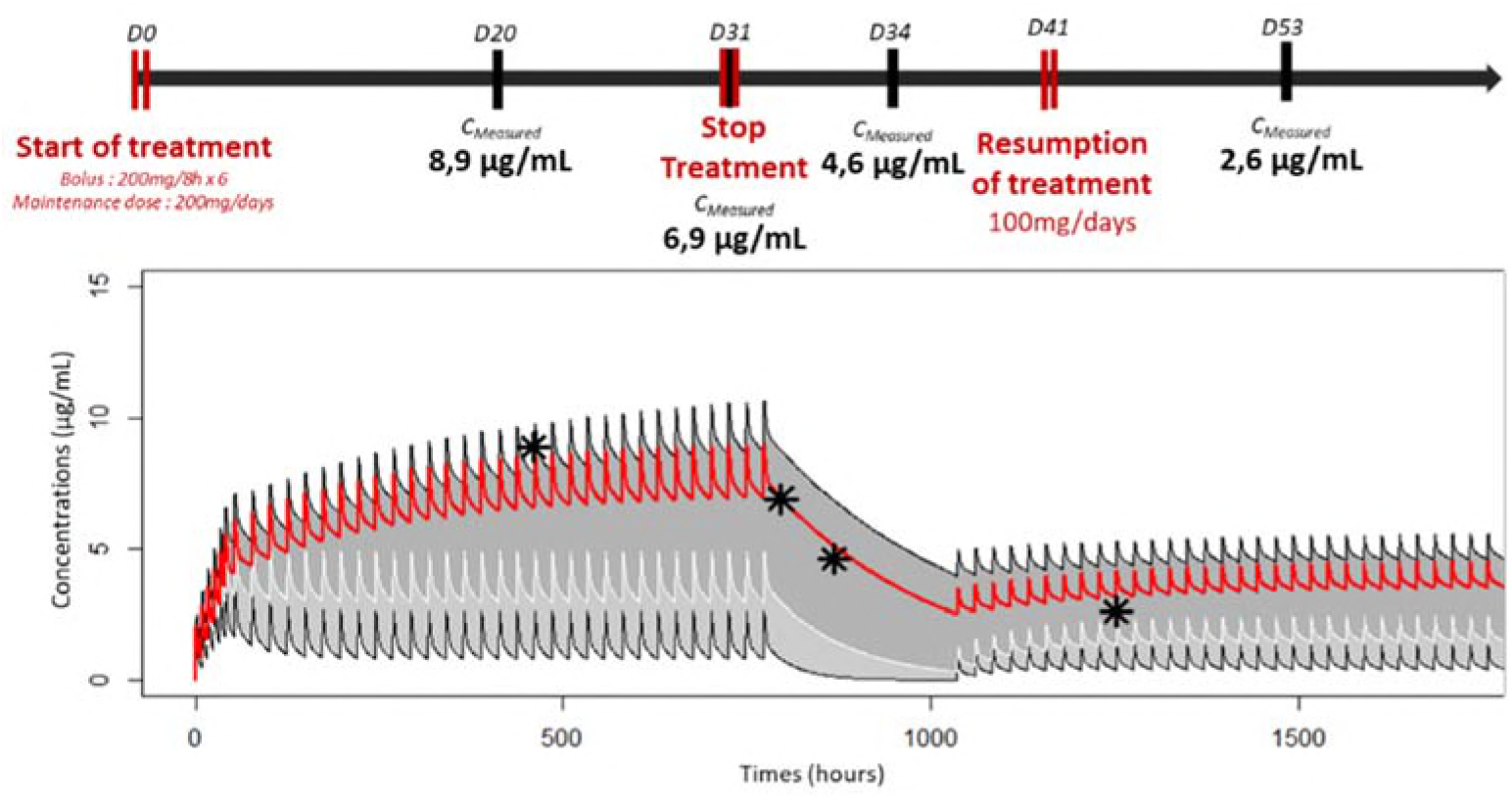
Patient 2: Kinetic profile of isavuconazole. Median kinetic profile (curve in white), “extremes” profiles found in less than 5% and 95% of the population (black curves) and estimated kinetic profile of the patient (red curve); the black stars represent the measured concentrations.

**Fig. 3.**
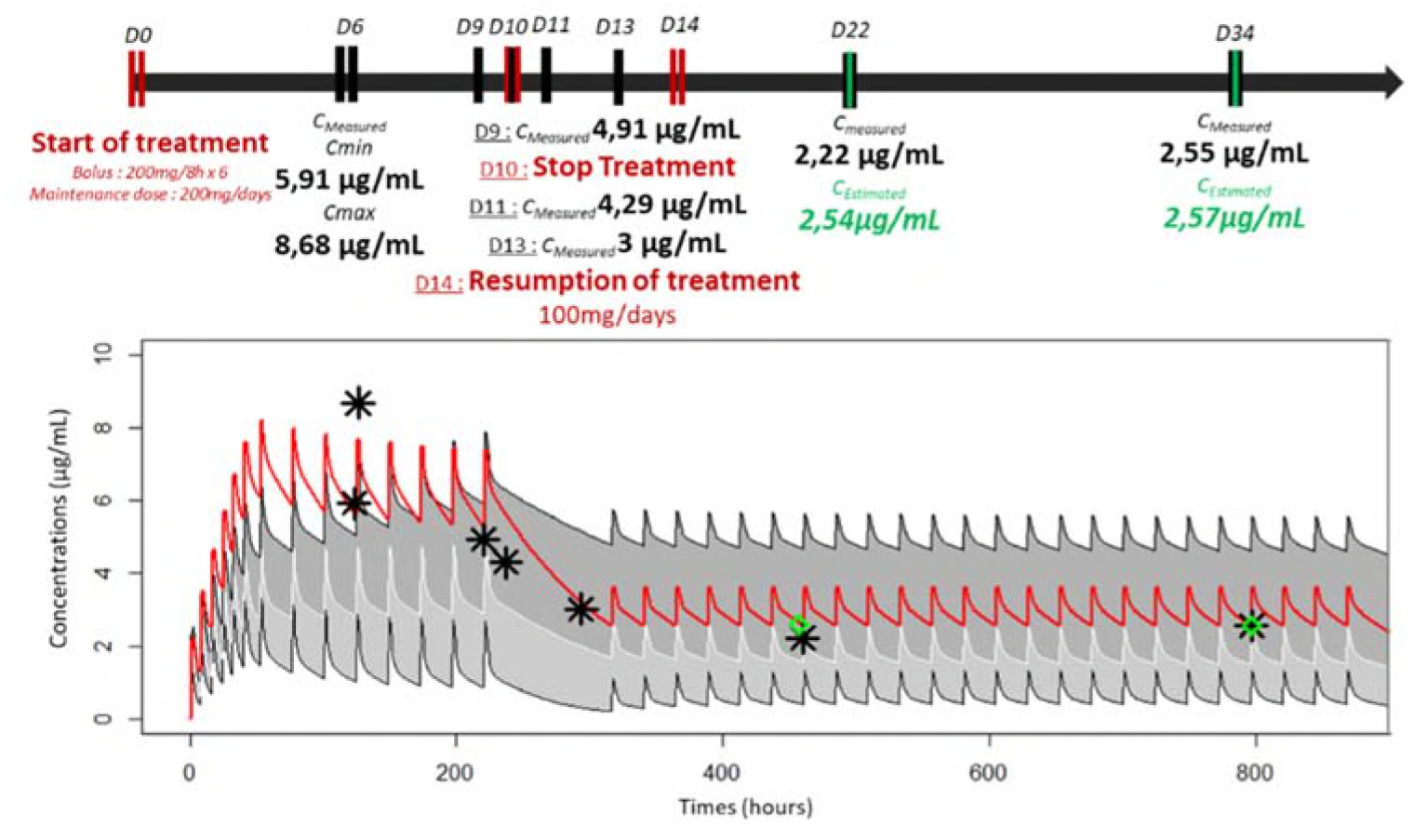
Patient 3: Kinetic profile of isavuconazole. Median kinetic profile (curve in white), “extremes” profiles found in less than 5% and 95% of the population (black curves) and estimated kinetic profile of the patient (red curve); the black stars represent the measured concentrations while the green ones represent the estimated trough concentrations in case of individualized dosage adjustment and the therapeutic drug monitoring confirmed the estimated values.

**Fig. 4.**
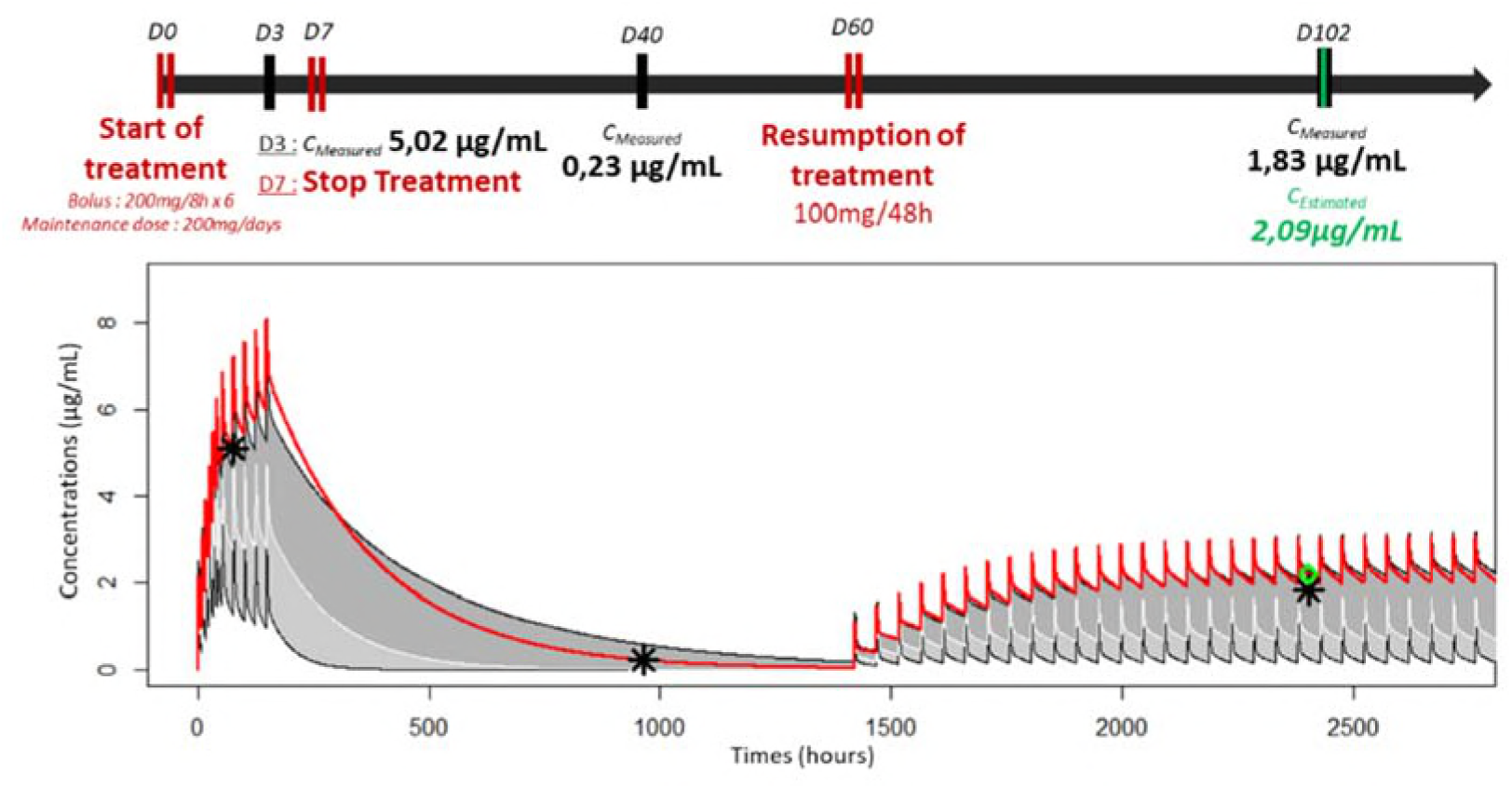
Patient 4: Kinetic profile of isavuconazole. Median kinetic profile (curve in white), “extremes” profiles found in less than 5% and 95% of the population (black curves) and estimated kinetic profile of the patient (red curve); the black stars represent the measured concentrations while the green ones represent the estimated trough concentrations in case of individualized dosage adjustment and the therapeutic drug monitoring confirmed the estimated values.

**Table 1.**
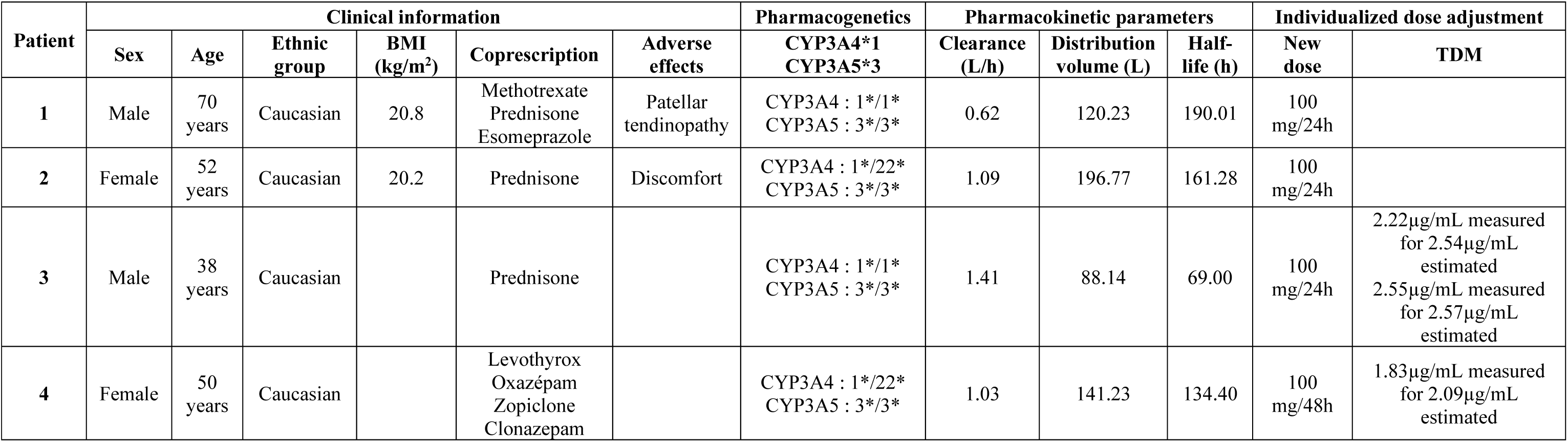
Pharmacokinetic-pharmacogenetic exploration of isavuconazole for 4 French patients presenting high unexpected plasma levels: this exploration was done as part of the hospital care of patients. *TDM* therapeutic drug monitoring Column 1: patient’s number Column 2 to 7: Clinical information found for each patient Column 8: Pharmacogenetic polymorphisms found for each patient according to a real-time PCR technique Column 9 to 11: Pharmacokinetic parameters of isavuconazole estimated from Desai’s population pharmacokinetic model [4] and plasma concentrations measured at different times depending on each patient Column 12 to 13: The estimated trough concentrations in case of individualized dosage adjustment

As isavuconazole is mainly metabolized by hepatic CYP3A4 and 3A5 [5–6], a pharmacogenetic exploration of *CYP3A4*22* (rs35599367) and *CYP3A5*3* (rs776746) polymorphisms, likely to explain low isavuconazole metabolism [7], was performed in these patients by real-time PCR.

This pharmacokinetic-pharmacogenetic exploration was routinely done as part of the hospital care of patients according to the regulations applied in France (patient’s informed consent for genotyping).

For all patients, pharmacokinetic parameters were different from the median values (clearance 2.36 L/h, peripherical distribution volume 241.28 L, elimination half-life 90.57 h) reported by Desai [4] and in particular the clearance that was systematically lower (Table 1). In accordance with clinicians, an individualized dose adjustment could be proposed for two patients (Fig.3 and 4, Table 1) and the therapeutic drug monitoring confirmed in each case that the estimated values were very closed to the measured ones (2.54 μg/mL and 2.57 μg/mL vs. 2.22 μg/mL and 2.55 μg/mL respectively for patient 3; 2.09 μg/mL vs. 1.83 μg/mL for patient 4). Among the 4 patients studied, all of them carried the homozygous variant *CYP3A5*3*, associated to a lake of CYP3A5 activity, while 2 patients were homozygous for the *CYP3A4*22*. In the Caucasian population, that is close to the Desai’s study population (83.2% of Caucasian’s and 16.8% of Asian’s patients [4]), more than 80-90% of patients possess *CYP3A5*3* polymorphism [8]. Consequently, this information cannot explain over-exposure in our patients. On the contrary, *CYP3A4*22* polymorphism is present in less than 8-10% of the Caucasian population [9]. Only 2 patients (50% of our population, patient 2 and 4) presented this polymorphism suggesting that other(s) explaining factor(s) is(are) likely elsewhere.

Based on these unexpected high concentrations and as previously suggested by Stott et al [1], we propose two recommendations. The first one consists in collecting a blood sample just before the first maintenance dose to estimate precociously the patient’s likely kinetic profile using Desai’s population pharmacokinetic model. Based on this profile, an individualized dose adjustment can be proposed as necessary. The second recommendation consists in screening CYP3A4 and 3A5 genetic polymorphisms, besides those usually explored (*CYP3A4*22* and *CYP3A5*3*), including the nonfunctional alleles *CYP3A4*17* (rs4987161) [10], *CYP3A5*6* (rs10264272) [8] and *CYP3A5*7* (rs76293380) [8] for patients presenting unexpected kinetic profiles. To date for patients in real life, too much sparse information concerning factors affecting pharmacokinetic profiles is available.

## ACKNOWLEDGEMENTS

The authors did not receive any funding for this project: the data have been generated as part of the routine work.

Léa Darnaud received grants from Gilead outside the submitted work. Cendrine Godet received consultancy or speaker fees, travel support from Pfizer, Astellas, Gilead, Basilea, MSD, SOS Oxygene and ISIS Medical outside the submitted work. Sandrine Pontier received grants from Astrazeneca and Novartis outside the submitted work. Alexia Debard received grants from Janssen-Cilag and Gilead outside the submitted work. Nicolas Venisse received grants from Gilead and Janssen-Cilag outside the submitted work. Peggy Gandia has received grants from Gilead and MSD outside the submitted work. All other authors: none to declare.

## REFERENCES

1. Stott KE and Hope WW. 2017. Therapeutic drug monitoring for invasive mould infections and disease: pharmacokinetics and pharmacodynamics considerations. Journal of Antimicrobial Chemotherapy. 72 (Suppl 1): i12–i18

2. Maertens JA, Raad II, Marr KA, Patterson TF, Kontoyiannis DP, Cornely OA, Bow EJ, Rahav G, Neofytos D, Aoun M, Baddley JW, Giladi M, Heinz WJ, Herbrecht R, Hope W, Karthaus M, Lee DG, Lortholary O, Morrison VA, Oren I, Selleslag D, Shoham S, Thompson GR, Lee M, Maher RM, Schmitt-Hoffmann AH, Zeiher B and Ullmann AJ. 2016. Isavuconazole versus voriconazole for primary treatment of invasive mould disease caused by Aspergillus and other filamentous fungi (SECURE): a phase 3, randomized-controlled, non-inferiority trial. The Lancet. 387: 20–26 (760-769)

3. Marty FM, Ostrosky-Zeichner L, Cornely OA, Mullane KM, Perfect JR, Thompson GR, Alangaden GJ, Brown JM, Fredricks DN, Heinz WJ, Herbrecht R, Klimko N, Klyasova G, Maertens JA, Melinkeri SR, Oren I, Pappas PG, Ráčil Z, Rahav G, Santos R, Schwartz S, Vehreschild JJ, Young JH, Chetchotisakd P, Jaruratanasirikul S, Kanj SS, Engelhardt M, Kaufhold A, Ito M, Lee M, Sasse C, Maher RM, Zeiher B and Vehreschild MJGT. 2016. Isavuconazole treatment for mucormycosis: a single-arm open-label trial and case-control analysis. The Lancet Infectious Diseases. 16 (7): 828–837

4. Desai A, Kovanda L, Kowalski D, Lu Q, Townsend R, and Bonate PL. 2016. Population pharmacokinetics of isavuconazole from phase 1 and phase 3 (SECURE) trials in adults and target attainment in patients with invasive infections due to aspergillus and other filamentous fungi. Antimicrob Agents Chemother. 60 5483–5491

5. Amsden JR and Gubbins PO. 2017. Pharmacogenomics of triazole antifungal agents: implications for safety, tolerability and efficacy. Expert Opinion on Drug Metabolism & Toxicology. 13 (11): 1135–1146

6. Townsend R, Dietz A, Hale C, Akhtar S, Kowalski D, Lademacher C, Lasseter K, Pearlman H, Rammelsberg D, Schmitt-Hoffmann A, Yamazaki T and Desai A. 2017. Pharmacokinetic evaluation of CYP3A4-mediated drug-drug interaction of isavuconazole with rifampin, ketoconazole, midazolam, and ethinyl estradiol/norethindrone in healthy adults. Clinical Pharmacology in Drug Development. 6 (10): 44–53

7. Gijsen VM, Schaik RH, Elens L, Soldin OP, Soldin SJ, Koren G and Wildt SN. 2013. CYP3A4*22 and CYP3A combined genotypes both corelate with tacrolimus disposition in pediatric heart transplant recipients. Pharmacogenomics. 14 (9): 1027–1036

8. Lamba J, Hebert JM, Schuetz EG, Klein TE and Altmanc RB. 2012. PharmGKB summary: very important pharmacogene information for CYP3A5. Pharmacogenetics and genomics. 22 (7): 555–558

9. Elens L, van Gelder T, Hesselink DA, Haufroid V and van Schaik RH. 2013. CYP3A4*22: promising newly identified CYP3A4 variant allele for personalizing pharmacotherapy. Review. Pharmacogenomics. 14 (1): 47–62

10. Dai D, Tang J, Rose R, Hodgson E, Bienstock RJ, Mohrenweiser HW and Goldstein JA. 2001. Identification of variants of CYP3A4 and characterization of their abilities to metabolize testosterone and chlorpyrifos. The Journal of pharmacology and experimental therapeutics. 299 (3): 825–831

